# A single-cell resolution gene expression atlas of the larval zebrafish brain

**DOI:** 10.1101/2022.02.11.479024

**Authors:** Inbal Shainer, Enrico Kuehn, Eva Laurell, Mariam Al Kassar, Nouwar Mokayes, Shachar Sherman, Johannes Larsch, Michael Kunst, Herwig Baier

**Affiliations:** Max Planck Institute of Neurobiology, Martinsried. Current address: Max Planck Institute for Biological Intelligence, in foundation, Martinsried, Germany; Allen Institute for Brain Science, Seattle, WA 98109, USA

**Author notes:** Co-first author (Inbal Shainer, Enrico Kuehn).

## Abstract

The advent of multimodal brain atlases promises to accelerate discoveries in neuroscience by offering in silico queries of cell types, connectivity and gene expression in regions of interest. We employed multiplexed fluorescent in situ RNA hybridization chain reaction (HCR) to generate expression maps for an initial set of 200 marker genes across the larval zebrafish brain. The data were registered to the Max Planck Zebrafish Brain (mapzebrain) atlas, thus allowing co-visualization of gene expression patterns, single-neuron tracings, transgenic lines and anatomical segmentations. Additionally, brain activity maps of freely swimming larvae were generated at single-cell resolution using HCR labeling of the immediate-early gene *cfos* and integrated into the atlas. As a proof of concept, we identified a novel class of cerebellar eurydendroid cells that express *calb2a*, project to the hypothalamus and are activated in animals that have recently ingested food. Thus, a cellular-resolution gene expression atlas will not only help with rapid identification of marker genes for neuronal populations of interest, but also bridge the molecular and circuit levels by anchoring genetic information to functional activity maps and synaptic connectivity.

## Introduction

Development and function of neural circuits depend on the spatiotemporal dynamics of differential gene expression across the brain. Particular classes of genes play prominent roles in organizing the layout of the nervous system and may serve as markers of cell-type identity and/or function. These include genes whose protein products directly influence cell-cell interactions and local properties of the synaptic network, and those that regulate the expression of other genes to determine cell fate, diversification, morphology and connectivity of neurons. Often the corresponding RNAs are expressed in a region- and cell-type specific fashion. Another class of important marker genes encode functional determinants, e. g., neurotransmitter-synthesizing enzymes, ion channels, synaptic machinery components, and neuropeptides; the presence of the corresponding proteins provides information about physiological properties of the expressing neurons and tends to be not cell-type specific (except for neuropeptides). Visualizing the expression of these markers in an atlas format offers a glimpse into the molecular and cellular composition of neural circuitry. Such an atlas then allows researchers to select markers for labeling and manipulating neurons of interest. Moreover, co-visualizing many of these markers offers a holistic view of the genetic architecture of the brain and may reveal general organizing principles.

The most prominent attempt at visualizing brain-wide transcript expression, while preserving spatial layout of the tissue, is probably the Allen Brain Atlas project, which published the results of RNA in situ hybridizations for thousands of genes in tissue sections, first in mouse (Lein et al., 2007) and later in human (Hawrylycz et al., 2012). These pioneering studies, together with both prior and more recent work with similar aims (e. g., Gong et al., 2003; Ortiz et al., 2020; M. Zhang et al., 2021), are of immense value for the research community. The resulting resources are routinely consulted by experimenters seeking to corroborate hypotheses and identify genetic markers for neuronal subpopulations of interest.

Atlases of the nervous system have also been devised for other model organisms, although, to our knowledge, not yet for RNA in situ hybridization patterns to faithfully map the endogenous gene expression. The nervous system of the nematode worm *Caenorhabditis elegans* has been charted in unsurpassed detail (Cook et al., 2019). Most recently, a strain of *C. elegans* has been developed by genetic engineering in which each of its 302 neurons is identifiable in one animal by a combination of fluorescent proteins (Yemini et al., 2021). This multi-color atlas enables in vivo interrogation of the spatial arrangement of identifiable neurons and serves as a convenient readout of the effects of developmental perturbations on cell fate and positioning (Yemini et al., 2021). In the fruitfly *Drosophila melanogaster*, the “virtual fly brain browser” (Milyaev et al., 2012) and the “fruit fly brain observatory” (Ukani et al., 2019) offer standardized neuroanatomical frames in which thousands of transgene expression patterns are co-registered, generating a light-microscopic mesoscale connectome. In the zebrafish *Danio rerio*, histological brain atlases have been available for some time (adult: Kenney et al., 2021, Wullimann et al., 1996; embryo/larva: Mueller and Wullimann, 2015), as well as atlas databases of hundreds of transgenic lines and immunostainings, such as “ViBE-Z” (Ronneberger et al., 2012), “Z-brain” (Randlett et al., 2015) and the “Zebrafish Brain Browser” (Tabor et al., 2019). The Max Planck Zebrafish Brain (“mapzebrain”) atlas (Kunst et al., 2019) at https://fishatlas.neuro.mpg.de/ is a multimodal digital atlas of the 6 days post-fertilization (6 dpf) larval brain, integrating data portals for transgenic lines, histological stains, and expertly curated neuroanatomical regions, as well as a growing number (currently ca. 4,300) of single-cell morphologies. These datasets are all registered into the same spatial coordinate system, the so-called ‘standard brain’ (Kunst et al., 2019). The atlas web interface offers refined visualization and animation tools, as well as options for downloading original data for offline analyses. Here we set out to add RNA fluorescent in situ hybridization (FISH) patterns to the mapzebrain atlas. Single-cell resolution was achieved by hybridization chain reaction (HCR), a spatial transcriptomics method that is highly sensitive, specific and multiplexable, allowing visualization of several RNA species in one specimen with different probe-specific fluorophores (Choi et al., 2018). Moreover, as a powerful tool for recording activity hotspots across the brain, we have devised an HCR FISH protocol for detecting *cfos* RNA following various sensory stimulations and behavioral tasks. All the data are publicly available and free to download through the mapzebrain website (https://fishatlas.neuro.mpg.de/). This new resource, which can be easily expanded in the future by community contributions, offers convenient web-based access to markers of neuronal subpopulations and paves the way for a genetic analysis of circuit architecture in this widely used vertebrate model.

## Results and Discussion

### Pipeline for HCR FISH labeling and registration to the mapzebrain atlas

Recent developments of FISH techniques, such as RNAscope, MERFISH and HCR (Chen et al., 2015; Choi et al., 2018; Wang et al., 2012), offer single RNA molecule detection and multiplexing in vivo. We chose HCR, because the staining protocol is easily implemented in larval and juvenile fish (Choi et al., 2018; Kappel et al., 2021) and is suitable for registration into a common 3D reference (Kappel et al., 2021; Lovett-Barron et al., 2020). HCR technology enables multiplexed FISH of up to 5 probe sets per round of staining with different fluorophores (Choi et al., 2018). To scale up the number of RNA species mapped, we tested two approaches: either multi-round HCR FISH by stripping the HCR signal by DNAse I treatment and re-staining of the same sample (Lovett-Barron et al., 2020; Xu et al., 2020); or labeling of different genes in different specimens and computational registering into a common coordinate system. Although the former strategy gives theoretically unlimited information about co-expression in specific single cells, we decided against it mainly for two reasons: First, we noticed that iterative rounds of stripping and re-labeling lead to a cumulative deformation of the tissue, which made alignment of the volume across staining rounds computationally challenging and prone to error. Second, the initial set of markers is, of course, not final, as new markers or better probes are certain to become available in the future. Thus, to allow for seamless addition of new data to the atlas, as well as for future improvements to the protocol, we chose to generate expression maps from single rounds of HCR and computational co-registration into a standard brain.

To enable co-alignment of different brains, we carried out HCR FISH stainings in the background of the *Tg(elavl3:H2b-GCaMP6s)* transgenic line, in which nuclear-localized GCaMP6s is expressed in almost all neurons. We generated an averaged brain template of the *Tg(elavl3:H2b-GCaMP6s)* line using the advanced normalization tool (ANTs) toolkit (Avants et al., 2011; Suppl. Figure 1) and calibrated a registration protocol (see Methods; Suppl. Movie 1). We then acquired FISH data for more than 200 genes (Suppl. Table 1), which were previously identified as distinct, often cell-type specific markers in retina, tectum and diencephalon (Kölsch et al., 2021; Raj et al., 2020; Sherman et al., 2021; Shainer et al., manuscript in preparation). This initial set of markers includes genes encoding transcription factors, cell-surface and secreted molecules, enzymes involved in transmitter synthesis and transport, neuropeptides, calcium-binding proteins, and other factors (Figure 1; Suppl. Table 1).

**Figure 1.**
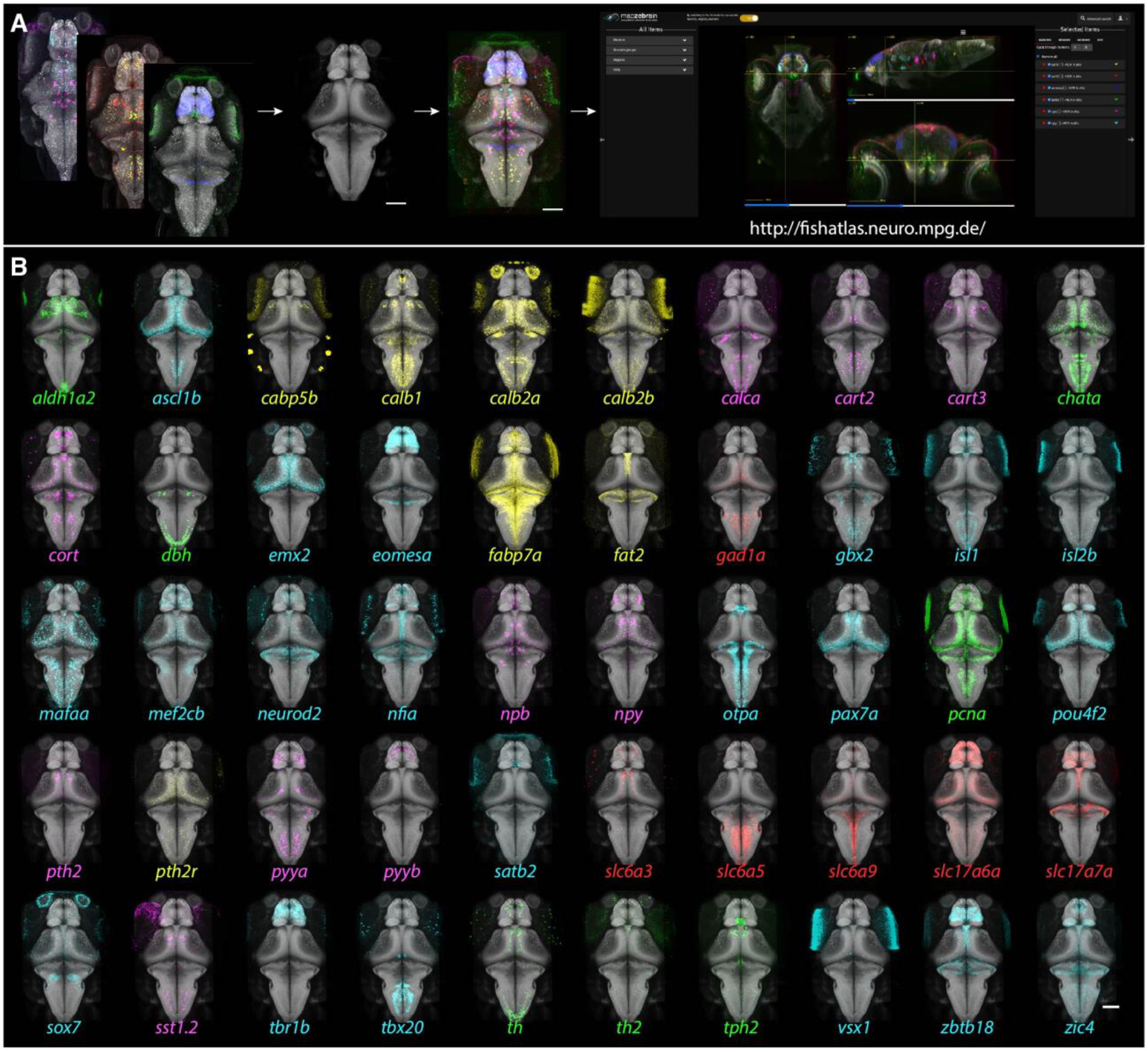
Gene expression atlas of the larval zebrafish brain. **A**. Multiplexed FISHs were performed on 6 dpf *Tg(elavl3:H2b-GCaMP6s)* fish. The GCaMP6s signal was utilized to align the images to the reference brain, and the aligned imaged were then uploaded to the mapzebrain atlas. Scale bar = 100 µm. **B**. 3D projections of 50 selected FISH images, registered into the standard brain. Color coding: green – enzymes; cyan – transcription factors; red – neurotransmitter transporters; magenta – neuropeptides; yellow – miscellaneous other. Scale bar = 100 µm.

For each marker gene, we labeled and imaged three 6 dpf fish, which we registered to the standard brain and averaged (Figure 1; see Methods). We multiplexed two genes per sample (Suppl. Table 1), and acquired the data at single-cell resolution, revealing the number of cells expressing a particular gene per region of interest (ROI), as well as any overlapping expression (Figure 2). All The FISH patterns of individual marker genes, including the average and the individual fish, are publicly accessible through the latest version of mapzebrain at https://fishatlas.neuro.mpg.de/ (Figure 1).

**Figure 2.**
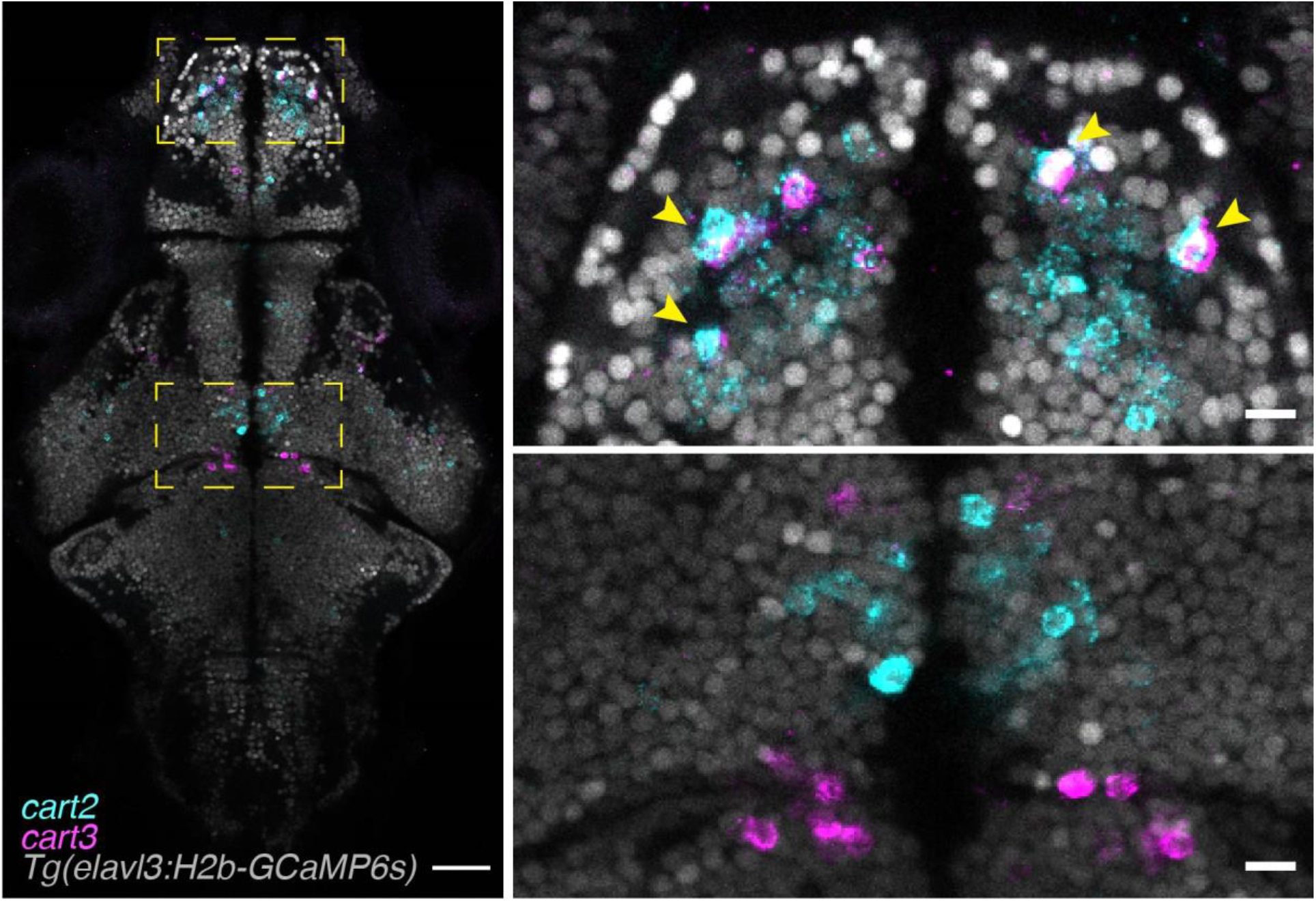
Multiplexed HCR in situ labeling of single cells. Single focal plane of a multiplexed HCR labeled fish. The endogenous expression of *Tg(elavl3:H2b-GCaMP6s)* labels neuronal nuclei and can be used to segment single neurons and to identify whether, or not, genes are co-expressed (arrowheads). Left scale bar = 100 µm, right scale bars = 10 µm.

### FISH image co-registration with other mapzebrain data modalities

Co-registration of the new FISH dataset with existing mapzebrain datasets frequently suggested spatial overlaps. For example, after registration, the expression of *aldh1a2* partially overlapped with the pattern of the transgenic enhancer trap line *mpn321Gt* in the pretectum (Suppl. Figure 2). To verify this in silico overlap of data acquired by different procedures (i. e., live imaging, immunostaining and FISH), we carried out *aldh1a2* FISH labeling in larvae expressing the *mpn321Gt* transgene and indeed found co-expression of the two markers in individual cells in the pretectum (Suppl. Figure 2). This finding highlights the potential to identify and confirm marker genes for cell populations of interest via rapid FISH labeling, and demonstrates the robustness and reproducibility of our digital image registration pipeline.

**Supplementary Figure 1.**
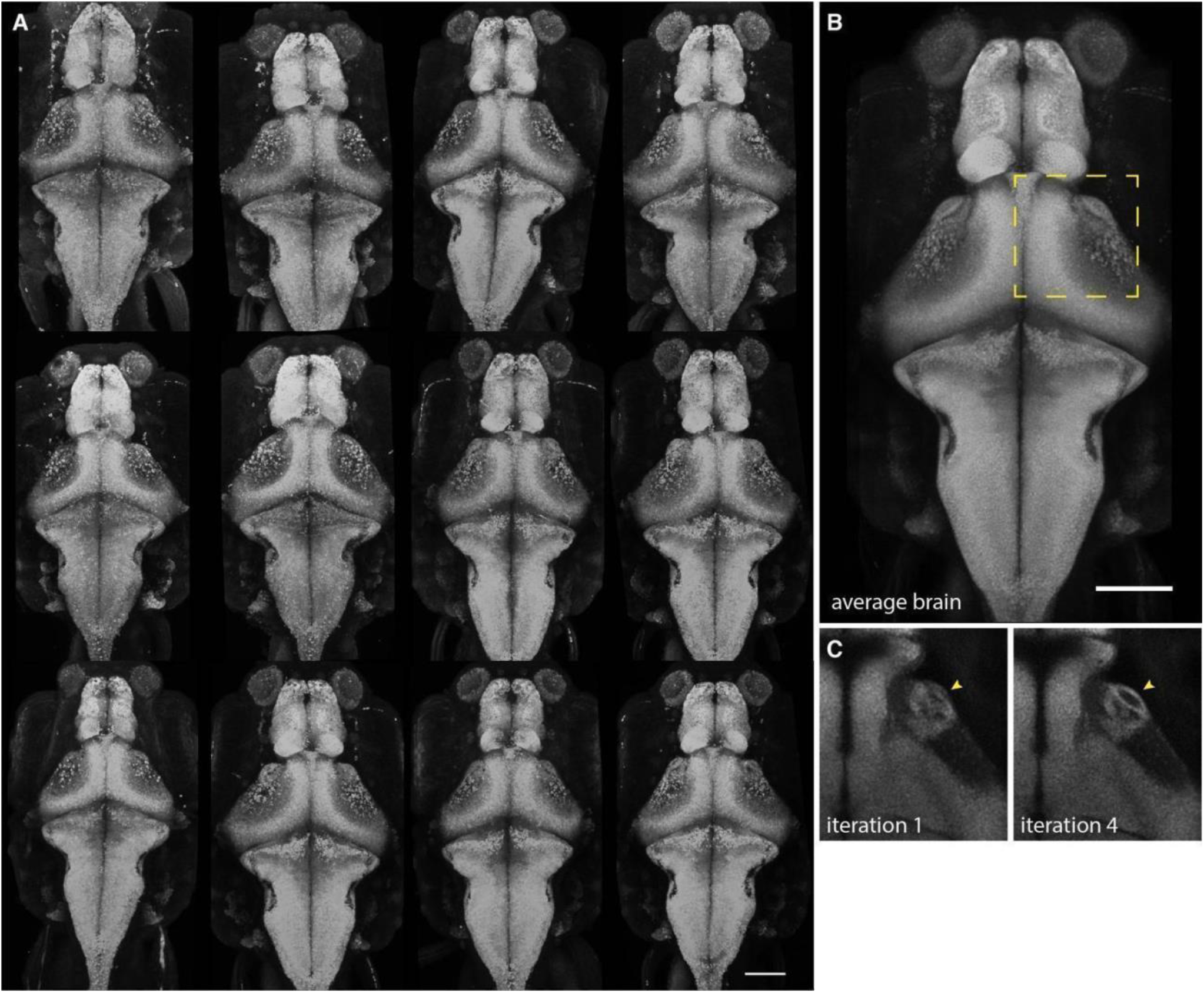
Generation of the FISH reference brain. **A**. 3D projections of 12 *Tg(elavl3:H2b-GCaMP6s)* age-matched (6 dpf) larvae. **B**. An unbiased average template was generated based on the individual images using ANTs iterative template construction. **C**. Template resolution was markedly improved over the 4 iteration rounds, as can, for example, be observed by the clear borders of arborization field 7 (arrowheads). Scale bars = 100 µm. **Suppl. Movie 1. Accurate registration of FISH images into the mapzebrain atlas**. An example of a single FISH sample after registration into the standard brain. Single focal planes of the GCaMP6s channel, spanning different depths (z planes) from ventral to dorsal, are shown. Scale bar = 100 µm.

**Supplementary Figure 2.**
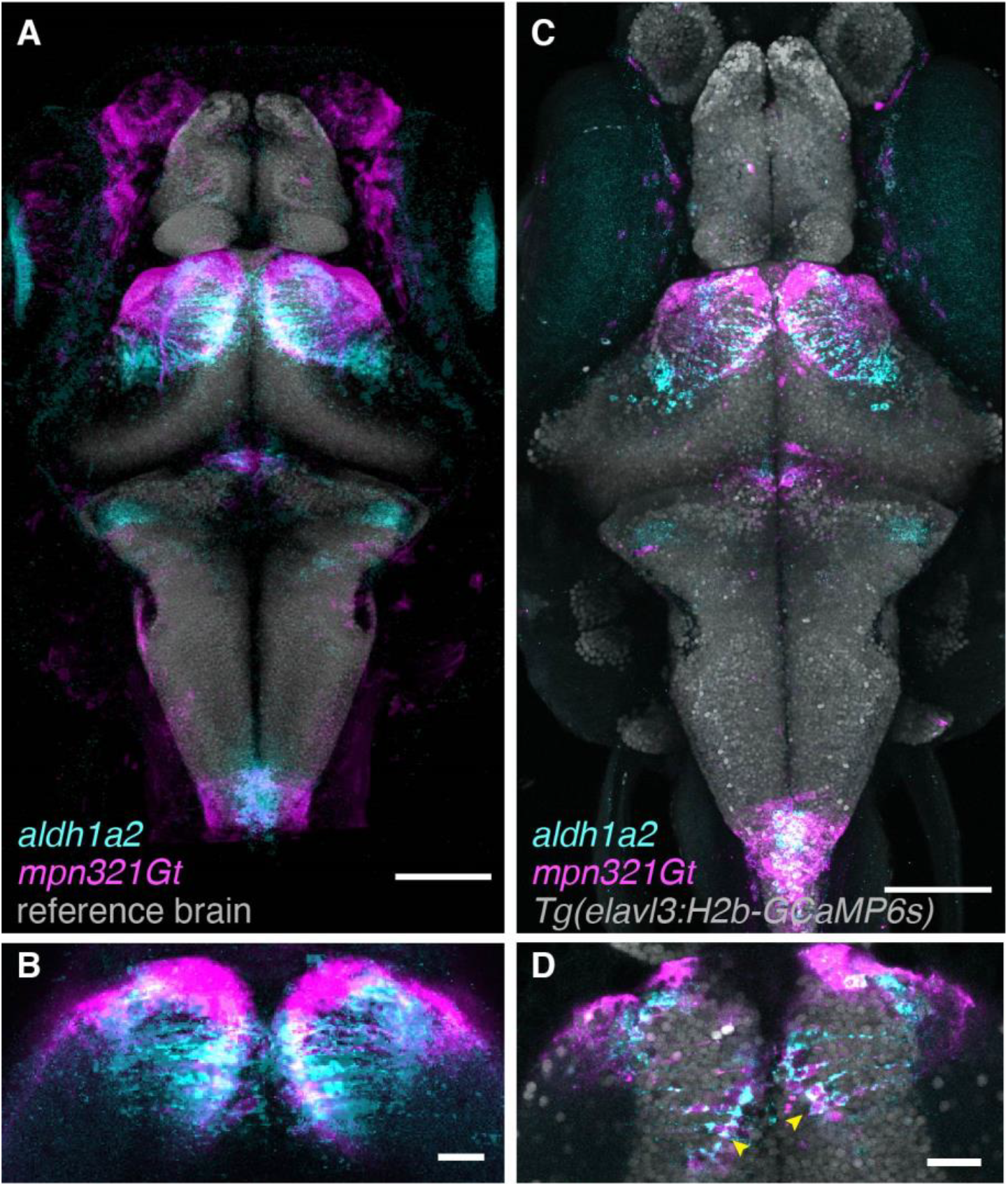
Registration correspondence between FISH and transgenic lines. **A**. 3D projection of *aldh1a2* FISH and *mpn321Gt* transgenic line after registration to the atlas. Scale bar = 100 µm. **B**. Single focal plane of the pretectum region of A. shows an in silico overlap between *aldh1a2* and *mpn321Gt*. Scale bar = 25 µm. **C**. 3D projection of *aldh1a2* FISH performed on a larva expressing both *mpn321Gt* and *Tg(elavl3:H2b-GCaMP6s)* transgenes. Scale bar = 100 µm. **D**. Single focal plane of the pretectum region of C. shows that *aldh1a2* is truly co-expressed together with *mpn321Gt* transgene in pretectum cells (arrowheads). Scale bar = 25 µm.

### Generating *cfos* brain activity maps of freely swimming fish in diverse behavioral paradigms

To expand the capabilities of our gene expression resource, we challenged the fish larvae in sensory or behavioral paradigms and carried out FISH staining of *cfos* (Figure 3; Suppl. Movie 2). *cfos* is an immediate early gene, expressed in neurons after prolonged depolarization, and is commonly used as a marker for neuronal activity (Herrera and Robertson, 1996; Kovacs, 2008). We exposed two groups of larval zebrafish to high temperature (32°C) or low temperature (22°C). Another group was allowed to hunt prey for 40 minutes. Other larvae were held for 40 minutes on a black, or a white, background. Still others were subjected for 40 min to a drifting, black-and-white grating, which induced an optomotor response (OMR; Neuhauss et al., 1999; Orger et al., 2000), or to a repeatedly looming black disk on a white background, thus evoking a series of visual escape responses (Temizer et al., 2015). Two groups were exposed either to an asymmetric light source, which evoked phototaxis (Brockerhoff et al., 1995; Orger et al., 2004), or to the alarm substance, ‘Schreckstoff’, which was extracted from fish skin (see Methods).

**Figure 3.**
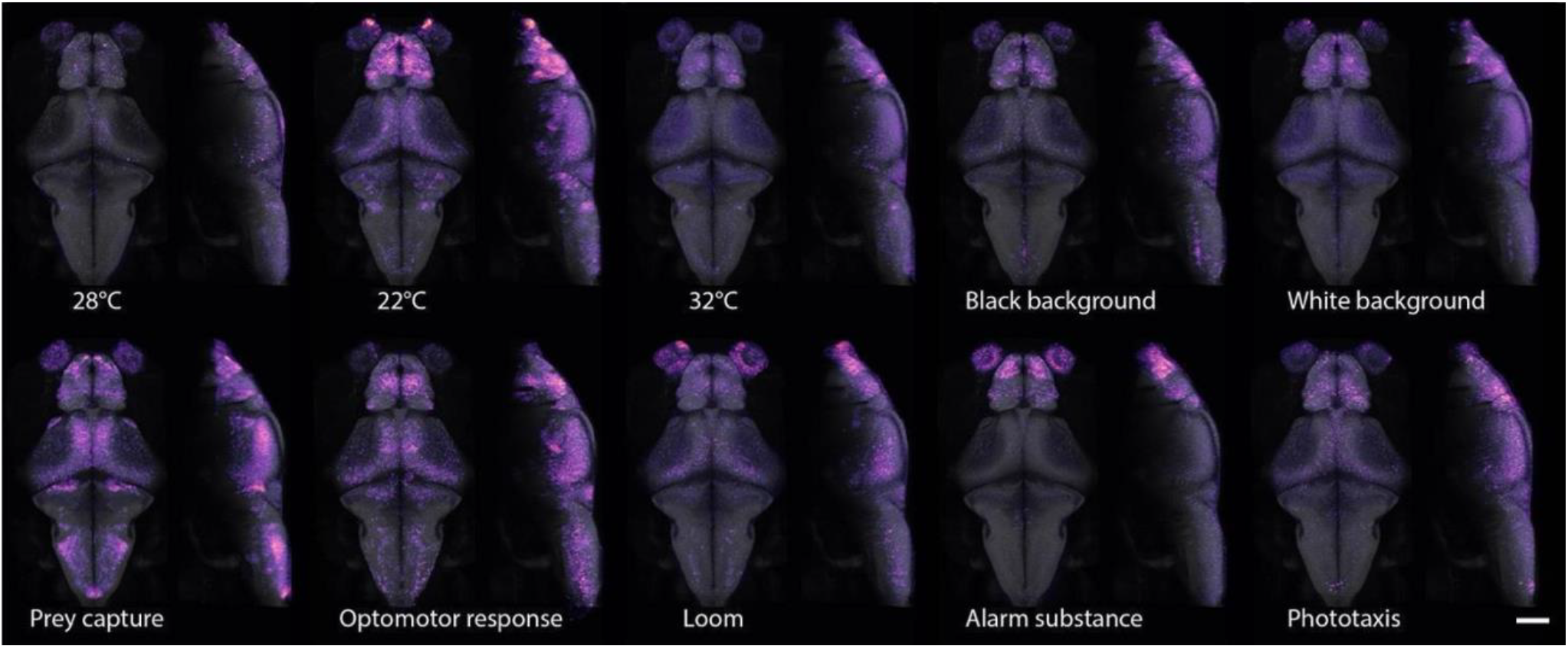
*cfos* brain activity maps of freely swimming fish. 3D projections of *cfos* FISH labeled areas of neuronal activity during behavior. Dorsal (left) and lateral (right) views are shown for each experimental condition. For interactive viewing, see mapzebrain atlas at https://fishatlas.neuro.mpg.de/. Scale bar = 100 µm. **Suppl. Movie 2. Brain activity maps of freely swimming fish**. Single focal planes spanning different depths (z planes) from ventral to dorsal. Several key anatomical areas are labeled. Scale bar = 100 µm. For interactive viewing, see mapzebrain atlas at https://fishatlas.neuro.mpg.de/.

For each condition, we imaged, registered and averaged six fish larvae. Qualitative inspection of the *cfos* FISH-labeled areas revealed different activity hotspots (Figure 3; Suppl. Movie 2). Activity maps largely confirmed previously published imaging studies of embedded animals (Chen et al., 2018; Chia et al., 2019; Semmelhack et al., 2014; Temizer et al., 2015; Wee et al., 2019). For example, in larvae that were hunting prey, we observed strong *cfos* labeling in the anterior optic tectum and retinal arborization field 7 (AF7; Figure 3; Suppl. Movie 2), which were extensively described for their involvement in prey detection and prey capture initiation in both embedded and freely swimming fish (Antinucci et al., 2019; Cong et al., 2017; Gahtan et al., 2005; Semmelhack et al., 2014; Z. Zhang et al., 2021).

Additionally, our approach revealed activity hotspots, which had not been reported previously. For example, in larvae that were hunting prey, we observed *cfos* labeling in cells surrounding the cerebellar neuropil (Figure 3; Suppl. Movie 2). Activity in this area seems to be specific to situations in which the animal is allowed to freely hunt and appears in conjunction with ingestion of food (Cong et al., 2017; Z. Zhang et al., 2021). We conclude that single-cell resolution *cfos* maps can help to localize brain activity in the context of unrestrained behavior.

### Discovery of behaviorally relevant circuits using *cfos* FISH

To explore how the *cfos* activity maps can be used for circuit analysis, we closely investigated the cerebellar area activated during hunting behavior. Interrogation of the mapzebrain atlas for marker genes expressed in this location, revealed that this region contains neurons labeled by *calb2a*. We verified this overlap by multiplexed HCR FISH of *cfos* and *calb2a* (Figure 4).

**Figure 4.**
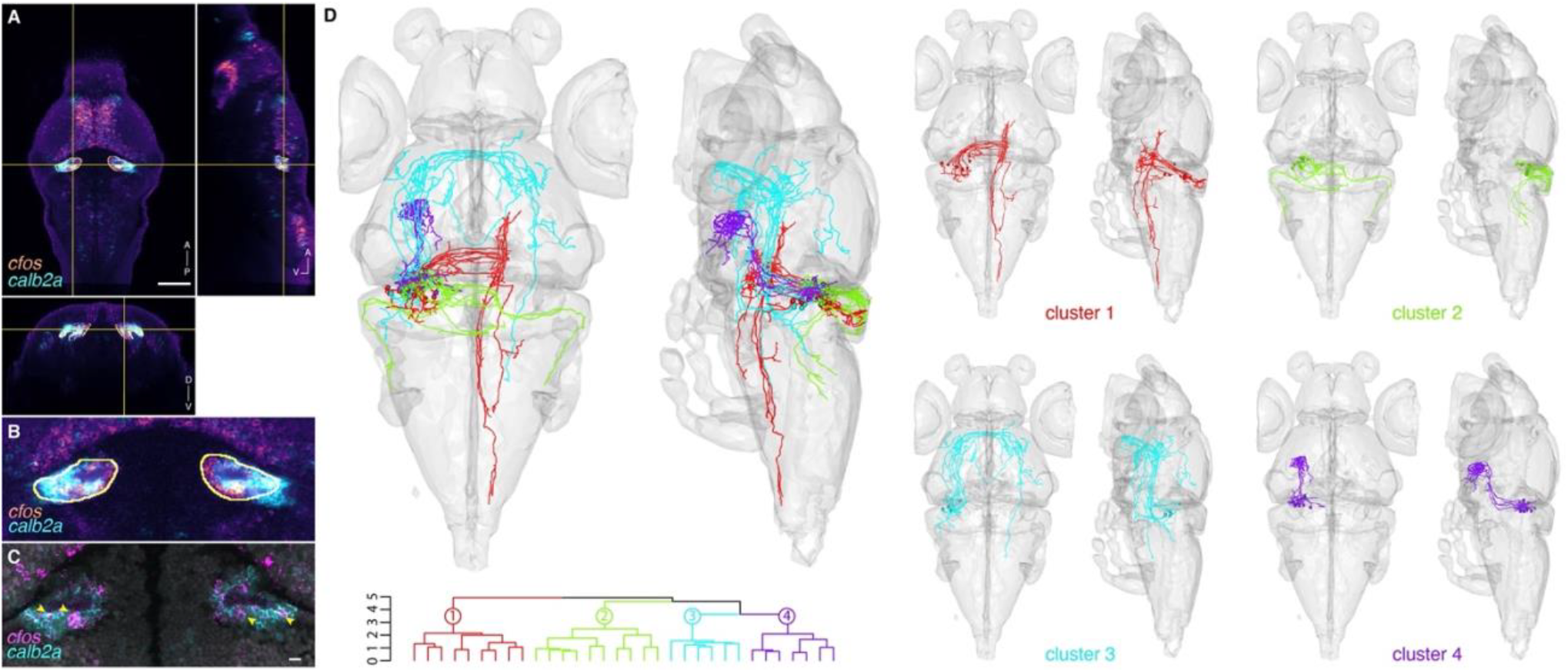
Marker gene expression and hypothetical connectivity of cerebellum neurons involved in prey capture. **A**. Orthogonal views of the *cfos* prey capture brain activity map with the registered *calb2a* FISH. An ROI is drawn around the *cfos* positive cerebellar area (yellow). Scale bar = 100 µm. Abbreviations: A-anterior; D-dorsal; P-posterior, V-ventral. **B**. Enlargement of the *cfos*-positive cerebellar area. Possible co-labeling was identified between the registered *cfos* and *calb2a* data. **C**. Multiplexed FISH of *calb2a* and *cfos* verified the suggested co-labeling (arrowheads), and therefore revealing a marker gene for the cerebellar neurons involved in prey capture. A single focal plane of an individual larva is shown. Scale bar = 10 µm. **D**. The ROI drawn in A. was used to search for single neurons whose somas are within the ROI. 32 neurons were identified, mirrored onto the left hemisphere and hierarchically clustered according to their morphology. One cluster of Purkinje cells (cluster 2), was identified as well as 3 clusters of eurydendroid cells, with projections to the contralateral hindbrain (cluster 1), to the contralateral retinal arborization field 7 (AF7) and tectum through the postoptic commissure (cluster 3) and to the hypothalamus (cluster 4).

Next, we drew a 3D ROI (‘volume of interest’) encompassing the cerebellar *cfos*-labeled region to query the database for individual morphologically reconstructed neurons from the mapzebrain atlas. We focused on neurons that had their somas within the ROI (Figure 4). Our search yielded 32 neurons, which we hierarchically clustered using the NBLAST algorithm (Costa et al., 2016) into 4 morphological types: one with stereotypical Purkinje cell morphology projecting to the octavolateral nucleus, and 3 eurydendroid cell types (Figure 4D). One group of eurydendroid cells projected to the dorsal zone of the periventricular hypothalamus (Figure 4D, cluster 4), an area involved in satiety response (Wee et al., 2019). Strikingly, this brain region was also strongly stained for *cfos* expression during prey capture behavior (Suppl. Movie 2). We speculate that *calb2a*^+^ eurydendroid cells, which are activated immediately after ingestion, contribute to regulation of satiety, perhaps through connectivity to the hypothalamus. This circuit may be conserved: Eurydendroid cells are considered homologous to the deep cerebellar nuclei in mammals (Bae et al., 2009; Heap et al., 2013). In humans and mice, deep cerebellar nuclei are activated after ingestion of food and may signal satiety (Low et al., 2021). While an experimental test of this hypothesis is beyond the scope of the current work, the newly discovered correspondence illustrates the power of integrating gene expression, functional *cfos* labeling, transgene and single-cell reconstructions in a multimodal atlas resource.

## Conclusion

We have presented here an essential addition to the brain atlas resources for larval zebrafish: whole-brain, single cell-resolution gene expression data for an initial set of 200 markers, co-registered in a single standard brain volume with thousands of single-neuron tracings, hundreds of transgenic lines and over 100 expertly curated anatomical segmentations. We further show, as a proof of concept, that such multimodal datasets, in combination with *cfos* mRNA brain activity maps, enable the discovery of behaviorally relevant neuronal subpopulations and pathways. The web interface of mapzebrain gene expression resource offers advanced computational, visualization and downloading options and can be seamlessly expanded in the future by community contributions to the atlas.

## Methods

### Zebrafish husbandry and maintenance

Adult zebrafish of the TLN strain were kept at 28°C with a day/night cycle of 14/10 hours, pH of 7-7.5, and a conductivity of 600 µS. All the experiments were performed on *Tg(elavl3:H2b-GCaMP6s)* larvae produced by natural matings. Larvae were raised until 6 days post fertilization (dpf) at 28°C in Danieau’s solution in petri dishes. The larvae were treated with PTU starting 24 hours post fertilization to prevent pigmentation. For *cfos* experiments, *Tg(elavl3:H2b-GCaMP6s)* larvae in the TLN background, carrying the *mitfa* mutation and lacking melanin pigment, were used without PTU treatment. The animal experiments were performed under the regulations of the Max Planck Society and the regional government of Upper Bavaria (Regierung von Oberbayern), approved protocols: ROB-55.2-2532.Vet_02-20-11, ROB-55.2-2532.Vet_02-21-93, 55.2-ROB-2532.Vet_02-20-183, ROB-55.2-2532.Vet_02-19-16.

### HCR FISH staining

HCR reagents, including probes, hairpins and buffers, were purchased from Molecular Instruments (Los Angeles, California, USA). The staining was performed according to a modified protocol of the *“HCR RNA-FISH protocol for whole-mount zebrafish embryos and larvae”* provided by Molecular Instruments (Choi et al., 2018). *Tg(elavl3:H2b-GCaMP6s)* positive larvae were anesthetized in 1.5 mM tricaine and fixed with ice-cold 4% PFA/DPBS overnight at 4°C with gentle shaking. The following day, larvae were washed 3 times for 5 min with DPBST (1x Dulbecco’s PBS + 0.1% Tween-20) to stop fixation, followed by a short 10 min treatment with ice-cold 100% Methanol at −20°C to dehydrate and permeabilize the tissue samples. Next, rehydration was performed by serial washing of 50% MeOH/50% DPBST and 25% MeOH/75% DPBST for 5 min each and finally 5 × 5 min in DPBST. 10-12 larvae were transferred into a 1.5 ml Eppendorf tube and pre-hybridized with pre-warmed hybridization buffer for 30 min at 37°C. Probe solution was prepared by transferring 2 pmol of each HCR probe set (2 µl of 1 µM stock) to 500 µl of hybridization buffer at 37°C (for *cfos* experiments, 4 µl of 1 µM probe stock were used.) The hybridization buffer was replaced with probe solution, and the samples were incubated for 12-16 hours at 37°C with gentle shaking. To remove excess probe, larvae were washed 4 × 15 min with 500 µl of pre-warmed probe wash buffer at 37°C. Subsequently, larvae were washed 2 × 5min with 5x SSCT (5x sodium chloride sodium citrate + 0.1% Tween-20) buffer at room temperature. Next, pre-amplification was performed by incubating the samples in 500µl of amplification buffer for 30 min at room temperature. Separately, 30 pmol of hairpin h1 and 30 pmol of hairpin h2 were prepared by snap-cooling 10 µl of 3 µM stock by incubating the hairpins in 95°C for 90 seconds, and cooling down to room temperature in a dark environment. After cooling down for 30 min, hairpin solution was prepared by transferring the h1 and h2 hairpins to 500 µl amplification buffer. The pre-amplification buffer was removed, and the samples were incubated in the hairpin solution for 12-16 hours in the dark at room temperature. Excess hairpins were washed the next day 3 × 20 min using 5x SSCT at room temperature. Larvae were then long-term stored at 4°C in 5X SSCT until imaging.

### Imaging

Samples were embedded in 2% low-melting agarose in 1x DPBS (Dulbecco’s PBS) and imaged with a Zeiss LSM700 confocal scanning microscope (upright), equipped with a 20x water immersion objective. Z-stacks, composing 2 tiles, were taken and stitched to produce a final image with size of 1039 × 1931 pixel (463.97 × 862.29 µm, 1 µm in z).

### Generation of reference brain and image registration and averaging

All the image registrations and averaging procedures were performed using Advanced Normalization Tools (ANTs) (Avants et al., 2011), which were previously calibrated and used for generation of zebrafish average brain images (Kunst et al., 2019; Marquart et al., 2017). Twelve transgenic fish expressing the transgenes *Tg(elavl3:H2b-GCaMP6s)* and *Tg(elavl3:lynTag-RFP)* were fixed and underwent the entire HCR buffers procedure, but without the usage of any probes or hairpins, and imaged as described. The images were used to generate an unbiased average template brain by running the following ANTs command on the GWDG computing cluster:

~~~
antsMultivariateTemplateConstruction2.sh -d 3 -o ${outputPath}/T_ -i 4
-g 0.2 -j 32 -v 500 -c 2 -k 2 -w 1×1 -f 12×8×4×2 -s 4×3×2×1 -q
200×200×200×200 -n 0 -r 1 -l 1 -z ${target1} -z ${target2} -m CC[2] -t
SyN[0.05,6,0.5] ${inputPath}
~~~

To register the newly generated reference to the mapzebrain coordinate system, we first registered the older *Tg(elavl3:H2b-GCaMP6s)* average image that existed in mapzebrain (Kunst et al., 2019) onto the newly generated reference and applied the inverse transformation on the newly generated reference image to align to the mapzebrain coordinate system. First, the following ANTs command was used to transform the older mapzebrain *Tg(elavl3:H2b-GCaMP6s)* average to new reference:

~~~
antsRegistration -d 3 --float 1 -o [${output1},${output2}] --
interpolation WelchWindowedSinc --use-histogram-matching 0 -r
[${template},${input1},1] -t rigid[0.1] -m
MI[${template},${input1},1,32,Regular,0.25] -c [200×200×200×0,1e-8,10] --
shrink-factors 12×8×4×2 --smoothing-sigmas 4×3×2×1vox -t Affine[0.1] -m
MI[${template},${input1},1,32,Regular,0.25] -c [200×200×200×0,1e-8,10] --
shrink-factors 12×8×4×2 --smoothing-sigmas 4×3×2×1 -t SyN[0.05,6,0.5] -m
CC[${template},${input1},1,2] -c [200×200×200×200×10,1e-7,10] --shrink-
factors 12×8×4×2×1 --smoothing-sigmas 4×3×2×1×0
~~~

Secondly, inverse transformation was applied on the newly generated reference using the inverse Warp transformation file by running the following ANTs command:

~~~
antsApplyTransforms -d 3 -v 0 -- float -n WelchWindowedSinc -i ${input3}
-r ${template} -o ${output4} -t
${output1}1InverseWarp.nii.gz -t ${output1}0GenericAffine.mat
~~~

The newly registered reference brain (Supp. Fig. 1 and Suppl. Movie 1) is available to download through the mapzebrain webpage. All the imaged HCR in situ images were registered onto this new reference by registering the *Tg(elavl3:H2b-GCaMP6s)* image channel using the ANTs command:

~~~
antsRegistration -d 3 --float 1 -o [${output1},${output2}] --
interpolation WelchWindowedSinc --use-histogram-matching 0 -r
[${template},${input1},1] -t rigid[0.1] -m
MI[${template},${input1},1,32,Regular,0.25] -c [200×200×200×0,1e-8,10] --
shrink-factors 12×8×4×2 --smoothing-sigmas 4×3×2×1vox -t Affine[0.1] -m
MI[${template},${input1},1,32,Regular,0.25] -c [200×200×200×0,1e-8,10] --
shrink-factors 12×8×4×2 --smoothing-sigmas 4×3×2×1 -t SyN[0.05,6,0.5] -m
CC[${template},${input1},1,2] -c [200×200×200×200×10,1e-7,10] --shrink-
factors 12×8×4×2×1 --smoothing-sigmas 4×3×2×1×0,
~~~

followed by applying the transformation files on the HCR image channels using the ANTs command

~~~
antsApplyTransforms -d 3 -v 0 -- float -n WelchWindowedSinc -i ${input3}
-r ${template} -o ${output4} -t ${output1}1Warp.nii.gz -t
${output1}0GenericAffine.mat
~~~

Each HCR-labeled RNA was imaged from three different fish, and registered as described. The three images were then arithmetically averaged using ImageJ (Schneider et al., 2012). Both the average image and the individual registered images are accessible through the mapzebrain site.

### *cfos* FISH experiments

For *cfos* experiments, larvae were raised until 6 dpf under optimal conditions (14/10h light cycle, 28°C) in Danieau’s solution. At 6 dpf, the larvae were transferred into a 6-well plate, 10 larvae per well until 7 dpf, when they were exposed to different stimuli for 40 min, followed by fixation and HCR staining, as described above. Six fish of each condition were imaged and registered as described before, followed by arithmetical averaging with ImageJ (Schneider et al., 2012). For visualization purposes of the averaged images, the contrast was linearly enhanced. Suppl. Movie 2 anatomical legend was generated using QuPath (Bankhead et al., 2017).

#### Temperature adaptation

The 6-well plate containing the fish was placed into a water bath set either to 22°C or 32°C with similar background color for 40 min.

#### Background adaptation

The 6-well plate containing the fish was placed either on white or black background, at 28°C for 40 min.

#### Prey capture

Paramecia were added into the 6-well plate containing the larvae. Food was not limiting: Many paramecia were still uneaten after 40 min.

#### Phototaxis

The 6-well plate containing the fish was placed on half-white and half-black background (splitting the well background color equally), at 28°C for 40 min.

#### Looming

Larvae were exposed to black looming disc stimuli using a setup previously described (Larsch and Baier, 2018; Pantoja et al., 2020). Briefly, groups of 10 larvae were placed into a 10 cm watch glass containing Danieau’s solution and left for 2 hours to acclimate to room temperature and white projector illumination from below. For stimulation, expanding black discs were projected to the center of the dish from below. Discs expanded linearly from 0 - 10 cm diameter within 500 ms, once every 60 seconds. This cycle was repeated over 40 minutes.

#### Optomotor response

Larvae were exposed to moving gratings using a setup previously described (Larsch and Baier, 2018; Pantoja et al., 2020). Briefly, groups of 10 larvae were placed into a 10 cm watch glass containing Danieau’s solution and left for 2 hours to acclimate to a background of a stationary grating (black/white step grating, 10 mm spatial frequency) projected from below. After 2 hours, gratings were moved in alternating directions at 1 cycle per second in the following sequence: 15 s in one direction, 5 s pause, and 15 s in the opposite direction. This sequence was repeated over 40 minutes. Visual inspection confirmed a strong behavioral optomotor response under these stimulus conditions. A control group of larvae remained on stationary gratings the entire time.

#### Alarm substance response

Alarm substance is stored in the club cells of zebrafish epidermis and released upon skin damage. For the experiment, five adult wild-type donor fish were euthanized and scales from both flanks were collected with the blunt side of a sterile razor blade. The scales were washed off with sterile-filtered Danieau’s solution and collected in a Falcon tube, avoiding any contamination with blood. The total 20 ml isolate was vortexed, filtered through a Whatman paper, and placed on ice until use. 500 µl of the isolate was distributed evenly over the wells of the 6-well plate containing the larvae.

### Single neuron morphology clustering

An ROI surrounding the cerebellar *cfos* mRNA-labeled area was labeled using the ImageJ segmentation editor plugin. The ROI was then used to search through the mapzebrain platform for neurons whose somas were located within it. 32 neurons were identified, flipped to the left (as described in Kunst et al., 2019), and a morphology similarity score between them was calculated using the NBLAST algorithm (Costa et al., 2016) by applying the nblast_allbyall function implemented in the R package NeuroAnatomy Toolbox (nat) (Bates et al., 2020) with default parameters. Next, unsupervised hierarchical clustering was performed using the nhclust function of the nat package with default parameters, and the results were plotted using the R packages dendroextras and rgl. The analysis and plotting R code can be found at (https://github.com/ishainer/Shainer_Kuehn_et_al_2022).

## Funding

Alexander von Humboldt foundation research fellowship for postdoctoral researchers – I.S

NARSAD Young Investigator Award – J.L

Max Planck Society – H.B

## Acknowledgments

We thank Robert Kasper for microscopy consultation and support, and Krasimir Slanchev, Manuel Stemmer and Mario Wullimann for their input and support with experiments. We thank all members of the Baier lab for beta testing and fruitful discussions.

