## Supplementary figures and images for "A single-cell resolution gene expression atlas of the larval zebrafish brain"

### Supplementary Figure 1

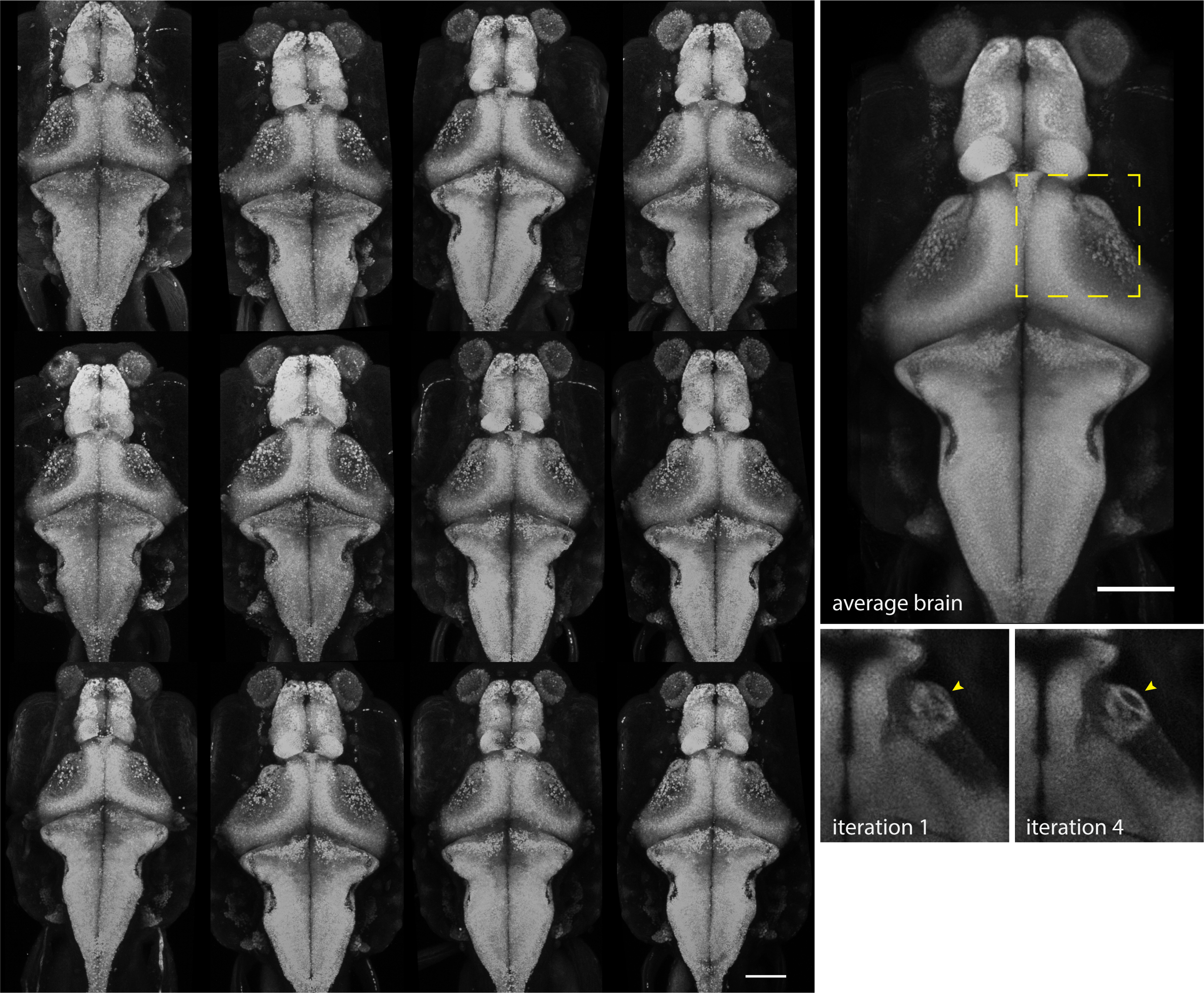

### Supplementary Figure 2

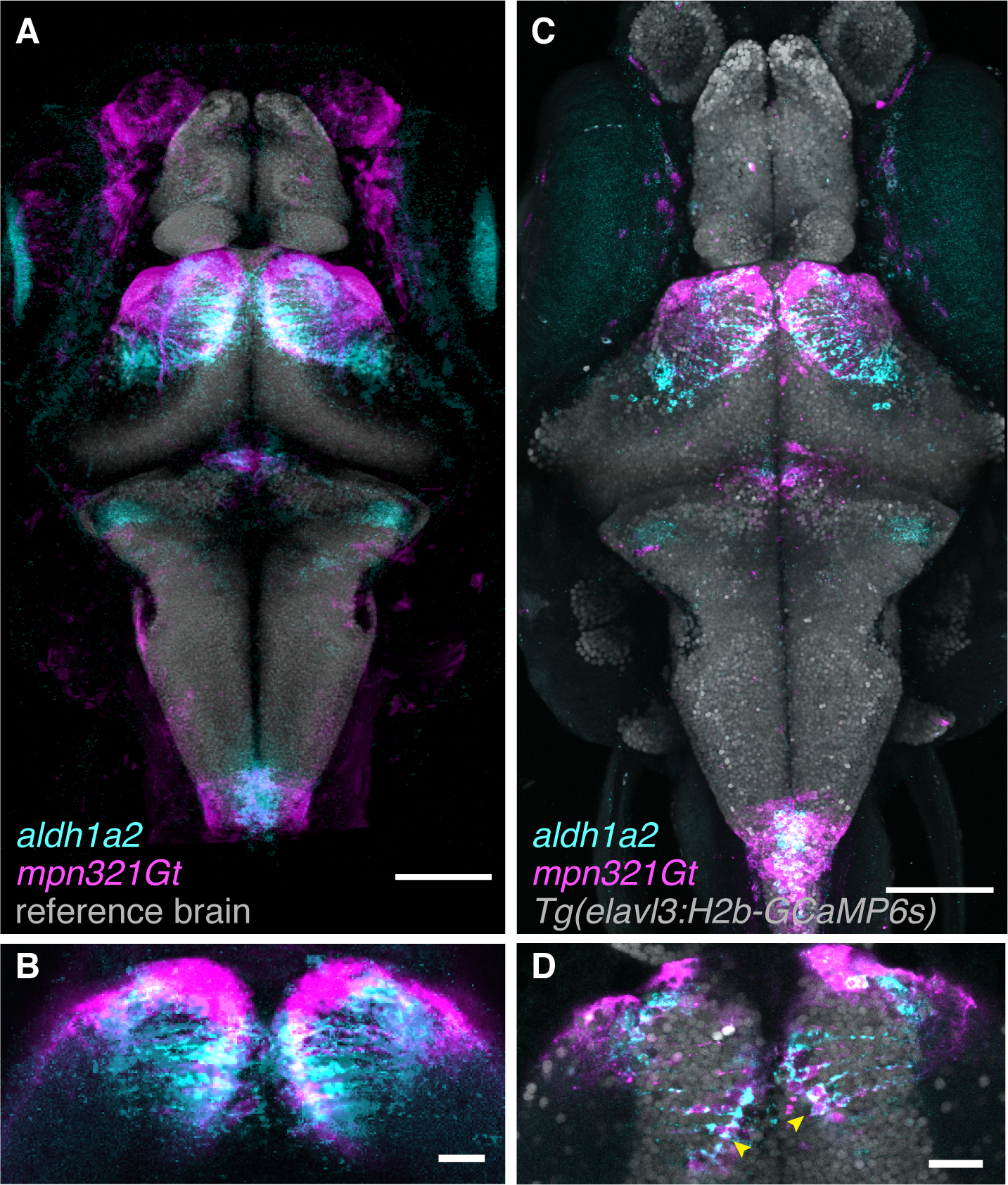
