## Supplementary table 1 for "A single-cell resolution gene expression atlas of the larval zebrafish brain"

| | gene name | NCBI refseq/ensembl ID | multiplexed with |
|---|---|---|---|
| 1 | *agrn* | NM_001177452.1 | *aldh1a2* |
| 2 | *agrp* | NM_001328012.1 | *asip2b* |
| 3 | *AL928650.3* | ENSDART00000152525.2 | *BX088596.1* |
| 4 | *aldh1a2* | NM_131850.1 | *agrn* |
| 5 | *ascl1a* | NM_131219.1 | *shha* |
| 6 | *ascl1b* | NM_131231.1 | *pou4f2* |
| 7 | *asip2b* | NM_001271291.1 | *agrp* |
| 8 | *atf5a* | NM_001128731.1 | *cebpa* |
| 9 | *atf5b* | ENSDART00000112838.3 | *nrgna* |
| 10 | *atoh7* | ENSDART00000101328.5 | *esrrga* |
| 11 | *avp* | NM_178293.2 | *calca* |
| 12 | *barhl1b* | ENSDART00000004548.8 | *fabp7a* |
| 13 | *bhlhe23* | NM_001030133.1 | *enc3* |
| 14 | *BX088596.1* | ENSDART00000173326.2 | *AL928650.3* |
| 15 | *c-fos (fosab)* | NM_205569.1 | - |
| 16 | *c1ql3b* | NM_205701.1 | *pnoca* |
| 17 | *cabp5b* | NM_001020732.1 | *calb1* |
| 18 | *CABZ01073265.1* | ENSDART00000162353.2 | *CR848683.2* |
| 19 | *calb1* | XM_001341568.5 | *cabp5b* |
| 20 | *calb2a* | NM_200718.1 | *calb2b* |
| 21 | *calb2b* | NM_200711.1 | *calb2a* |
| 22 | *calca* | NM_001002471.1 | *avp* |
| 23 | *cart2* | NM_001017570.1 | *cart3* |
| 24 | *cart3* | ENSDART00000052029.7 | *cart2* |
| 25 | *ccka* | XM_001346104.6 | - |
| 26 | *cckb* | ENSDART00000163899.2 | *rspo1* |
| 27 | *cebpa* | NM_131885.2 | *atf5a* |
| 28 | *chata* | NM_001130719.1 | *chatb* |
| 29 | *chatb* | KF840480.1 | *chata* |
| 30 | *chodl* | NM_001017578.2 | *npy* |
| 31 | *chrm4a* | ENSDART00000173727.2 | *chrm4b* |
| 32 | *chrm4b* | ENSDART00000157276.2 | *chrm4a* |
| 33 | *chrna2b* | XM_021468614.1 | *rpp25b* |
| 34 | *CR848683.2* | ENSDART00000194953.1 | *CABZ01073265.1* |
| 35 | *crabp1a* | NM_182858.1 | *nr5a1a* |
| 36 | *crema* | XM_677932.9 | *nocta* |
| 37 | *crh1b (crf, chrb)* | NM_001007379.1 | *crhbp* |
| 38 | *crha* | ENSDART00000136022.2 | *tcima* |
| 39 | *crhbp* | NM_001003459.1 | *crh1b (crf, chrb)* |
| 40 | *crx* | ENSDART00000037879.7 | - |
| 41 | *dacha* | NM_152955.2 | *gch1* |
| 42 | *dazap1* | NM_001173504.1 | *foxb1a* |
| 43 | *dbh (dopamine beta-hydroxylase)* | NM_001109694.2 | *slc6a3* |
| 44 | *dbx1a* | NM_131158.1 | *insm2* |
| 45 | *dlx5a* | NM_131306.2 | *lhx9* |
| 46 | *dmbx1a* | NM_178163.2 | *sp5l* |
| 47 | *drd4a* | ENSDART00000012603.9 | *ghrh* |

| | | | |
|---|---|---|---|
| 48 | *drgx* | NM_001037105.1 | *sst7 (cort)* |
| 49 | *emx2* | NM_131280.2 | *mafaa* |
| 50 | *en1b* | NM_001013498.2 | *evx1* |
| 51 | *enc3* | NM_001001402.2 | *bhlhe23* |
| 52 | *eomesa* | NM_131679.3 | *satb2* |
| 53 | *esrrb* | NM_001324539.1 | *pcbp3* |
| 54 | *esrrga* | NM_212954.1 | *atoh7* |
| 55 | *evx1* | NM_131249.2 | *en1b* |
| 56 | *fabp7a* | NM_131605.2 | *barhl1b* |
| 57 | *fat2* | XM_001920023.7 | - |
| 58 | *fat4* | XM_021481296.1 | *nfixb* |
| 59 | *fezf2* | NM_131636.2 | *otx2b* |
| 60 | *foxb1a* | NM_131285.1 | *dazap1* |
| 61 | *ftr82* | NM_001075103.1 | *tsnare1* |
| 62 | *gad1a* | XM_021478820.1 | - |
| 63 | *gad1b* | NM_194419.1 | *gad2* |
| 64 | *gad2* | NM_001017708.2 | *gad1b* |
| 65 | *galn* | NM_001346239.1 | *itpr1b* |
| 66 | *gbx2* | NM_152964.1 | *neurog1* |
| 67 | *gch1* | NM_001136255.2 | *dacha* |
| 68 | *gfra1a* | NM_131730.1 | *onecut1* |
| 69 | *ghrh* | NM_001080092.1 | *drd4a* |
| 70 | *gjd2b* | NM_194420.1 | *inhbaa* |
| 71 | *gnat1* | ENSDART00000064896.6 | *sema3fa* |
| 72 | *grm2b* | NM_001287547.1 | *nsun2* |
| 73 | *gyg1b* | NM_001002062.1 | *hcn1* |
| 74 | *hcn1* | ENSDART00000163878.2 | *gyg1b* |
| 75 | *her4.1* | NM_001103128.1 | *uncx* |
| 76 | *hpdb* | NM_001003742.1 | *rasd2* |
| 77 | *id2b* | NM_199541.1 | *six3b* |
| 78 | *inhbaa* | NM_130916.1 | *gjd2b* |
| 79 | *insm2* | XM_001332478.8 | *dbx1a* |
| 80 | *irx1b* | NM_131823.1 | *man2a2* |
| 81 | *isl1* | ENSDART00000010896.6 | *isl2b* |
| 82 | *isl2b* | ENSDART00000055936.5 | *isl1* |
| 83 | *itpr1b* | XM_021479879.1 | *galn* |
| 84 | *lhx9* | NM_001017710.2 | *dlx5a* |
| 85 | *mafaa* | NM_001082940.2 | *emx2* |
| 86 | *mafba* | NM_131015.3 | *mafbb* |
| 87 | *mafbb* | NM_131842.2 | *mafba* |
| 88 | *man2a2* | XM_009297800.2 | *irx1b* |
| 89 | *mc1r* | NM_180970.1 | *mc5rb* |
| 90 | *mc3r* | NM_180972.2 | *mc4r* |
| 91 | *mc4r* | NM_173278.1 | *mc3r* |
| 92 | *mc5ra* | NM_173279.1 | *pomca* |
| 93 | *mc5rb* | NM_173280.1 | *mc1r* |
| 94 | *mcm2* | ENSDART00000159629.3 | *mef2cb* |
| 95 | *mef2cb* | ENSDART00000135911.3 | *mcm2* |

| | | | |
|---|---|---|---|
| 96 | *nefma* | NM_001111214.2 | - |
| 97 | *neurod1* | NM_130978.2 | *neurod2* |
| 98 | *neurod2* | NM_131082.1 | *neurod1* |
| 99 | *neurod6a* | NM_131816.3 | *neurod6b* |
| 100 | *neurod6b* | NM_001309843.1 | *neurod6a* |
| 101 | *neurog1* | NM_131041.1 | *gbx2* |
| 102 | *nfia* | NM_001079962.1 | *pcna* |
| 103 | *nfil3-6* | NM_001002218.1 | *ntn1b* |
| 104 | *nfixb* | NM_001040248.1 | *fat4* |
| 105 | *ngb* | NM_131853.1 | *zic2a* |
| 106 | *nocta* | XM_695702.8 | *crema* |
| 107 | *npb* | NM_001127369.1 | - |
| 108 | *npy* | NM_131074.2 | *chodl* |
| 109 | *nr3c1 (GR)* | NM_001020711.3 | *nr3c2 (MR)* |
| 110 | *nr3c2 (MR)* | NM_001100403.1 | *nr3c1 (GR)* |
| 111 | *nr5a1a* | NM_131794.1 | *crabp1a* |
| 112 | *nrgna* | NM_001302620.1 | *atf5b* |
| 113 | *nsun2* | NM_199711.1 | *grm2b* |
| 114 | *ntn1b* | NM_130998.1 | *nfil3-6* |
| 115 | *olig3* | NM_001110393.1 | *olig4* |
| 116 | *olig4* | NM_199514.1 | *olig3* |
| 117 | *onecut1* | NM_200573.1 | *gfra1a* |
| 118 | *opn4b* | NM_001258224.1 | *opn4xa* |
| 119 | *opn4xa* | NM_001256077.1 | *opn4b* |
| 120 | *otpa* | NM_001128703.1 | *pcp4l1* |
| 121 | *otx2b* | NM_131251.1 | *fezf2* |
| 122 | *oxtra (oxtr)* | NM_001199370.1 | *oxtrb (oxtrl)* |
| 123 | *oxtrb (oxtrl)* | NM_001199369.1 | *oxtra (oxtr)* |
| 124 | *pax3a* | NM_131277.1 | *pax6a* |
| 125 | *pax6a* | NM_131304.1 | *pax3a* |
| 126 | *pax7a* | NM_131332.2 | *pax7b* |
| 127 | *pax7b* | NM_001146149.1 | *pax7a* |
| 128 | *pcbp3* | NM_001020731.2 | *esrrb* |
| 129 | *pcna* | ENSDART00000076304.5 | *nfia* |
| 130 | *pcp4l1* | NM_001161488.2 | *otpa* |
| 131 | *penkb* | NM_182883.1 | *r3hdm2* |
| 132 | *phex* | NM_001320330.1 | *rbpms2b* |
| 133 | *pitx2* | NM_130975.2 | - |
| 134 | *pmch* | NM_001202542.1 | *pmchl* |
| 135 | *pmchl* | NM_001162488.1 | *pmch* |
| 136 | *pnoca* | XM_003200329.5 | *c1ql3b* |
| 137 | *pnocb* | NM_001015044.1 | *sema3fb* |
| 138 | *pomca* | NM_181438.3 | *mc5ra* |
| 139 | *pou4f2* | NM_212807.1 | *ascl1b* |
| 140 | *ptf1a* | ENSDART00000021987.6 | *rx1* |
| 141 | *pth2* | NM_205577.3 | *pth2r* |
| 142 | *pth2r* | NM_131377.1 | *pth2* |
| 143 | *pyya* | NM_001164371.1 | *pyyb* |

| | | | |
|---|---|---|---|
| 144 | *pyyb* | NM_001327895.1 | *pyya* |
| 145 | *r3hdm2* | XM_021477336.1 | *penkb* |
| 146 | *rasd2* | NM_200532.1 | *hpdb* |
| 147 | *rbpms2b* | ENSDART00000006619.8 | *phex* |
| 148 | *rpp25b* | NM_001351699.1 | *chrna2b* |
| 149 | *rspo1* | NM_001002352.1 | *cckb* |
| 150 | *rx1* | ENSDART00000106166.5 | *ptf1a* |
| 151 | *satb1a* | ENSDART00000056205.7 | *satb1b* |
| 152 | *satb1b* | ENSDART00000160784.2 | *satb1a* |
| 153 | *satb2* | ENSDART00000088876.4 | *eomesa* |
| 154 | *scrt1b* | NM_001014347.2 | *sp9* |
| 155 | *sema3fa* | NM_001014822.1 | *gnat1* |
| 156 | *sema3fb* | XM_021479878.1 | *pnocb* |
| 157 | *shha* | NM_131063.3 | *ascl1a* |
| 158 | *si:dkey-237h12.3* | XM_021481239.1 | *znf536* |
| 159 | *six3b* | NM_131363.1 | *id2b* |
| 160 | *slc17a6a (vglut2b)* | NM_001009982.1 | *slc17a6b (vglut2a)* |
| 161 | *slc17a6b (vglut2a)* | NM_001128821.1 | *slc17a6a (vglut2b)* |
| 162 | *slc17a7a (vglut1)* | NM_001098755.1 | *slc17a7b (vglut1b)* |
| 163 | *slc17a7b (vglut1b)* | XM_009297642.3 | *slc17a7a (vglut1)* |
| 164 | *slc6a3 (dopamine transporter)* | NM_131755.1 | *dbh* |
| 165 | *slc6a5 (glyt2)* | NM_001009557.1 | *slc6a9 (glyt1)* |
| 166 | *slc6a9 (glyt1)* | NM_001030073.1 | *slc6a5 (glyt2)* |
| 167 | *sox14* | NM_001037680.1 | *sox1b* |
| 168 | *sox1b* | NM_001037662.1 | *sox14* |
| 169 | *sox7* | NM_001080750.2 | *tph2* |
| 170 | *sp5l* | NM_194371.2 | *dmbx1a* |
| 171 | *sp9* | NM_212960.2 | *scrt1b* |
| 172 | *sst1.1* | NM_183070.1 | *sst5* |
| 173 | *sst1.2* | NM_001386222.1 | *sstr1a* |
| 174 | *sst2* | NM_001128784.2 | *sst6* |
| 175 | *sst5* | XM_001333046.7 | *sst1.1* |
| 176 | *sst6* | NM_001128784.2 | *sst2* |
| 177 | *sst7 (cort)* | NM_001045431.2 | *drgx* |
| 178 | *sstr1a* | ENSDART00000159963.2 | *sst1.2* |
| 179 | *sulf2a* | NM_200936.1 | *tbx2b* |
| 180 | *tac1* | NM_001256391.1 | *tac3b* |
| 181 | *tac3b* | NM_001256390.1 | *tac1* |
| 182 | *tbr1b* | NM_001115090.3 | *tbx20* |
| 183 | *tbx20* | NM_131506.2 | *tbr1b* |
| 184 | *tbx2b* | ENSDART00000122101.4 | *sulf2a* |
| 185 | *tcima* | NM_001013472.2 | *crha* |
| 186 | *tfap2a* | NM_001319158.1 | *tfap2e* |
| 187 | *tfap2b* | NM_001024665.1 | *tfap2d* |
| 188 | *tfap2d* | NM_001025546.1 | *tfap2b* |
| 189 | *tfap2e* | NM_200821.2 | *tfap2a* |
| 190 | *th* | NM_131149.1 | *th2* |
| 191 | *th2* | NM_001001829.1 | *th* |

| | | | |
|---|---|---|---|
| 192 | *tph2* | NM_001310068.1 | *sox7* |
| 193 | *tsnare1* | XM_005158305.4 | *ftr82* |
| 194 | *txn* | NM_001002461.1 | *uts1* |
| 195 | *uncx* | NM_001020780.2 | *her4.1* |
| 196 | *uts1* | NM_001030180.1 | *txn* |
| 197 | *vsx1* | ENSDART00000078763.3 | *zbtb18* |
| 198 | *zbtb18* | NM_001082952.1 | *vsx1* |
| 199 | *zic1* | NM_130933.1 | *zic4* |
| 200 | *zic2a* | NM_131558.1 | *ngb* |
| 201 | *zic4* | NM_001076612.1 | *zic1* |
| 202 | *znf536* | XM_009303408.3 | *si:dkey-237h12.3* |
